# The ubiquitin ligase UBR-1 regulates the synaptic strength between the GABAergic and glutamatergic signaling

**DOI:** 10.1101/2023.08.26.554963

**Authors:** Yi Li, Jyothsna Chitturi, Bin Yu, Yongning Zhang, Jing Wu, Panpan Ti, Wesley Hung, Mei Zhen, Shangbang Gao

## Abstract

Excitation/Inhibition (E/I) balance is carefully maintained by the nervous system. Neurotransmitter GABA has been reported to be co-released with its sole precursor, another neurotransmitter glutamate. The genetic and circuitry mechanisms to establish the balance between GABAergic and Glutamatergic signaling have not fully elucidated. *C. elegans* DVB is a classically defined excitatory GABAergic motoneuron that drives the expulsion step in defecation motor program. We show that in addition to UNC-47, the vesicular GABA transporter, DVB also expresses EAT-4, a vesicular glutamate transporter. UBR-1, a conserved ubiquitin ligase, regulates the DVB activity by suppressing a bidirectional inhibitory glutamate signaling. Loss of UBR-1 impairs the DVB Ca^2+^ activity and the expulsion frequency. These impairments are fully compensated by the knock-down of EAT-4 in DVB. Further, glutamate-gated chloride channels GLC-3 and GLC-2/4 receive DVB’s glutamate signals to inhibit DVB and enteric muscle, respectively. These results implicate an intrinsic cellular mechanism that promotes the inherent asymmetric neural activity. We propose that the elevated glutamate in *ubr-1* being the cause of the E/I shift, potentially contributes to the Johanson Blizzard Syndrome.

## Introduction

A neural network maintains an equilibrium between excitatory (E) and inhibitory (I) synaptic signaling (Yizhar *et al*, 2011). Disruptions to the E/I balance, such as by increased or decreased levels of inhibitory signaling, appear to be an emerging theme in neurodevelopmental and neurological disorders (Rubenstein & Merzenich, 2003). Understanding the cellular processes that affect the E/I balance is critical to our understanding of neural homeostasis and associated disorders.

The ubiquitin-proteasome system facilitates the spatial and temporal regulation of protein metabolism and is thus involved in all aspects of cellular processes (Bachmair *et al*, 1986; Gardner *et al*, 2005; Haglund & Dikic, 2005; Varshavsky, 2014). A key component of the systems is the substrate-recognition factor, the ubiquitin ligases (E3s) (Hershko *et al*, 1983). UBR1 is an E3 ligase; loss-of-function mutations in human UBR1 cause the Johanson Blizzard Syndrome (JBS), a rare autosomal recessive disorder with broad developmental and neurological symptoms, including abnormal facial appearance, exocrine pancreatic insufficiency and varying degrees of mental retardation (Daentl *et al*, 1979; Johanson & Blizzard, 1971). Pathophysiology of JBS has remained unclear and there is currently no causal treatment (Sukalo *et al*, 2014). The yeast and mouse loss-of-function models for human UBR1’s closest homologs do not show JBS-like syndromes, or phenotypes that reveal leads to the underlying cellular defects underlying the JBS (Balogh *et al*, 2002; Varshavsky, 1996).

There has been considerable focuses on the identification of UBR1 substrates. Some critical substrates include the cohesion complex subunit SCC1, the transcriptional activator Msn4 and the hydroxyaspartate dehydratase Sry1 in yeast (Kim *et al*, 2014; Rao *et al*, 2001); the pluripotency factor LIN-28 in *C. elegans* (Weaver *et al*, 2017); the GTPase activators RGS4/5 (Hu *et al*, 2005; Lee *et al*, 2011), the breast cancer-associated tumor suppressor BRCA1 (Xu *et al*, 2012) and the PTEN induced putative kinase 1 PINK1 in mammalian cells (Yamano & Youle, 2013). However, whether these substrates share functional conservation across species, and how their dys-regulation might relate to the various JBS pathological features have not been established (Choi *et al*, 2010; Matta-Camacho *et al*, 2010).

An alternative and complementary approach to address the JBS is through characterization of genetic modifiers of phenotypes exhibited by an UBR1 animal model. This is of physiological relevance to reveal, for example, dys-regulated signaling pathways in the absence of UBR1, as genetic suppressors of altered behaviors (Kelly & Stanley, 2001).

*C. elegans* offers such an experimental platform (Chalfie *et al*, 1985; Gao *et al*, 2018; Huang *et al*, 2019; Kawano *et al*, 2011; Lim *et al*, 2016; Petrash *et al*, 2013; Stawicki *et al*, 2013). *C. elegans* has a single homologue of UBR family protein UBR-1 (**Fig EV1A**), which contains all main conserved functional domains and is consistently expressed in muscles and neurons (Chitturi *et al*, 2018). Previously, we have shown that the loss of UBR-1 alters glutamate homeostasis, leading to elevate glutamate level (Chitturi *et al*., 2018). One behavioral consequence of elevated glutamate is stiff body bending during backward movement caused by the simultaneous activation of the execution motor neurons (Chitturi *et al*., 2018).

Glutamate is an abundant and essential metabolite (Kelly & Stanley, 2001), as well as a predominant excitatory neurotransmitter. Being the sole precursor of γ-amino butyric acid (GABA), an inhibitory neurotransmitter (Costa *et al*, 1979), glutamate has been reported to be co-present with GABA in both glutamatergic and GABAergic neurons (Beltrán & Gutiérrez, 2012; Root *et al*, 2018; Vaaga *et al*, 2014). A rat model for depression exhibits reduced GABA/glutamate-mediated synaptic response ratio and increased GABA signaling after treatment with an antidepressant (Shabel *et al*, 2014). Whether elevated glutamate level in *C. elegans ubr-1* mutant leads to imbalanced GABA/glutamate signaling is an intriguing and unexplored question.

DVB exhibits rhythmic activity, contracting enteric muscles through an excitatory GABAergic signaling (Beg & Jorgensen, 2003; Branicky & Hekimi, 2006; Wang *et al*, 2013). In this study, we show that loss of UBR-1 impairs rhythmic expulsion, a motor behavior controlled by DVB. We show that DVB expresses not only the vesicular GABA transporter (vGAT) UNC-47 but also the vesicular glutamate transporter (vGluT) EAT-4. In *ubr-1* mutants, calcium (Ca^2+^) activity of the DVB neuron is decreased. The removal of EAT-4 from DVB fully rescued *ubr-1* mutant’s expulsion defect and restored the Ca^2+^ activity of DVB neuron. Removing either of the inhibitory glutamate gated chloride channels (GluCl), GLC-3 in DVB and GLC-2/GLC-4 in intestinal muscles also rescued the expulsion defects. Lastly, we demonstrate that exogenous glutamate perfusion led to an instantaneous inhibition of the Ca^2+^ activity of DVB, an effect dampened by low extracellular chloride and *glc-3* mutation. UBR-1, by gating the level of glutamate, maintains an inherent imbalance of excitatory and inhibitory signaling by the DVB neuron, specifically, by suppressing inhibitory glutamatergic signaling at a neuron functioning predominantly through excitatory GABAergic signaling.

## Results

### *ubr-1* loss-of-function mutants exhibit reduced expulsion in the defecation motor program

*ubr-1* mutants exhibit multiple motor defects. In addition to reduced body bending (Chitturi *et al*., 2018), they exhibit defect in defecation, a rhythmic motor program that occurs every 45-50 seconds.

The defecation motor program consists of a stereotypically ordered, three-step motor sequence: the posterior body contraction (pBoc), the anterior body contraction (aBoc), and the final expulsion step (Exp) (**Fig 1A**). We found that while the pBoc and aBoc steps in *ubr-1* were unaffected by *ubr-1* mutations (pBoc 13.6 ± 0.2/10 min, aBoc 13.6 ± 0.2/10 min in *ubr-1*; pBoc 13.23 ± 0.17/10 min, aBoc 13.23 ± 0.17/10 min in wild type) (**Fig 1B**), the expulsion frequency in *ubr-1(hp684)* loss-of-function mutants (7.55 ± 0.34/10 min) was significantly reduced when compared to wild-type animals (12.9 ± 0.21/10 min) (**Fig 1B**). This deficiency was fully rescued by restored expression of a UBR-1 genomic fragment under its endogenous promoter (P*ubr-1*) (**Fig 1B and C**). These observations suggest that the *ubr-1* gene is specifically required for the expulsion step.

**Figure 1.**
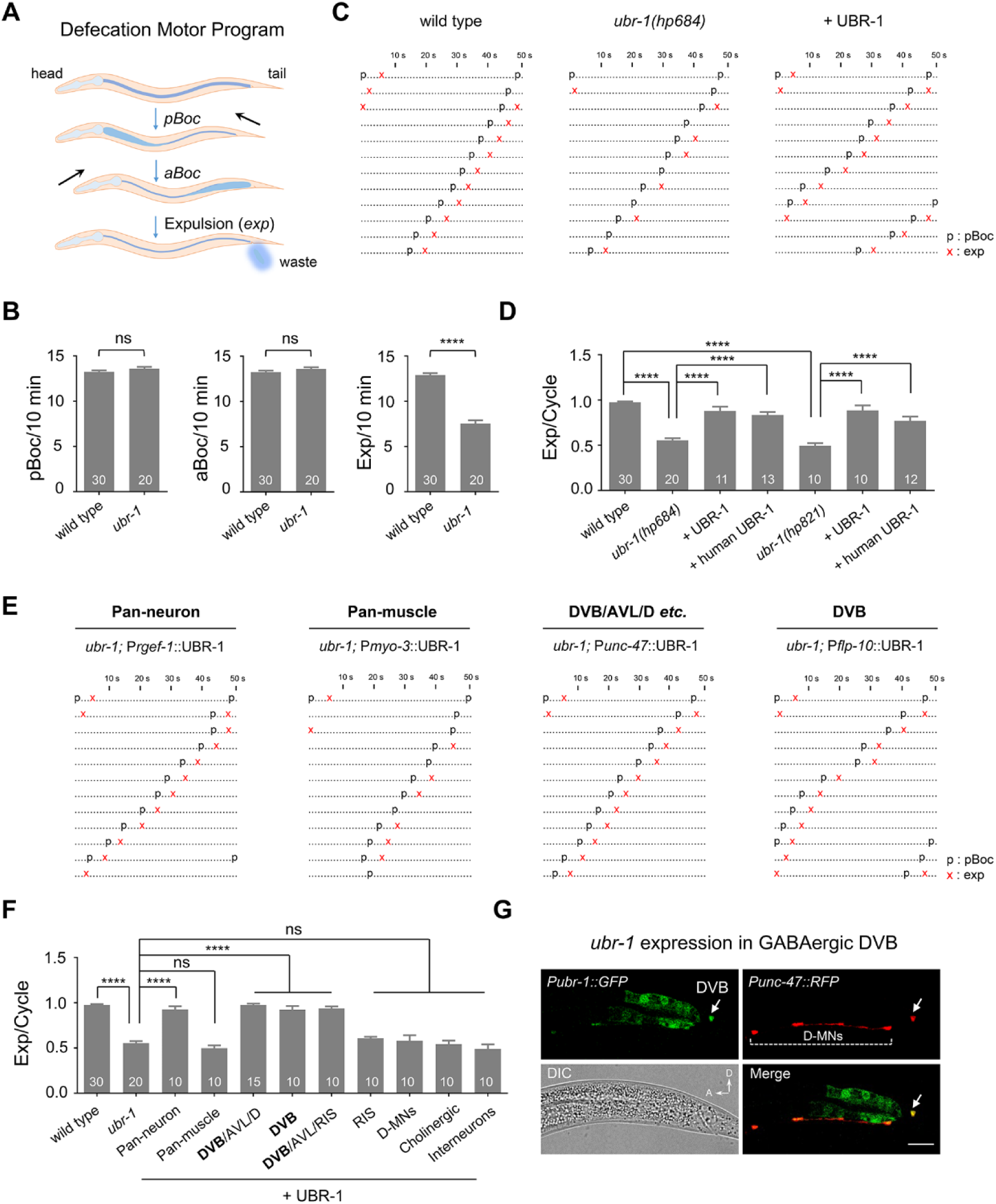
*ubr-1* regulates the defecation Exp step. (A) A schematic diagram of *C. elegans* defecation motor program (DMP). DMP is initiated by posterior body contraction (*pBoc*), and followed by anterior body contraction (*aBoc*) after ∼2 s relaxation phase and then enteric muscle contraction, leading to expulsion of the gut contents (*exp*). (B) Quantification of the frequency of pBoc, aBoc and Exp events in different genotypes. *ubr-1*(*hp684*) exhibits reduced expulsion frequency. \*\*\*\**p* < 0.0001; two-tailed unpaired *t*-test. (C) Representative ethograms of consecutive 10 min defecation cycles in wild type, *ubr-1* mutant and rescued worms. Each dot represents 1 s. ‘‘p’’ stands for pBoc and ‘‘x’’ indicates exp. aBoc is omitted. (D) Quantification of the expulsion rhythm in different *ubr-1* alleles. “Exp/Cycle” is specified as the ratio of Exp over pBoc. \*\*\*\**p* < 0.0001; one-way ANOVA test. (E, F) Representative ethograms of defecation cycles and quantification of the expulsion frequency in different genotypes. Reduced expulsion in *ubr-1* mutant could be rescued by restoring UBR-1 expression in DVB and AVL neurons (P*unc-47* and P*flp-10*) but not in GABAergic D-motor neurons (P*unc-25s*), cholinergic motor neurons (P*acr-2*), GLR-positive interneurons (P*glr-1*) or muscles (P*myo-3*). \*\*\*\**p* < 0.0001; one-way ANOVA test. (G) *ubr-1* expression in DVB neuron. White arrow denotes the DVB soma, and D-motor neurons are labeled. Scale bar, 20 µm. n = 10-30 animals. Error bars, SEM.

The *hp684* mutant allele leads to a nonsense truncation of the last 194 amino acids of UBR-1(Q1864X) (**Fig EV1B**), raising concerns of the residual protein function. We further analyzed the expulsion rate in three other *ubr-1* alleles that miss one or more critical conserved domains (**Fig EV1B**): *hp821*(E34X), which contains an N-terminal stop codon mutant that predicated the loss of all domains; *hp821 hp833* (E34X, E1315X), which contains an addition premature internal stop codon in the RING finger domain; and *hp865*, which lacks the entire RING finger domain by replacing it with an SL2-NLS::GFP (Chitturi *et al*., 2018). All alleles exhibited a similar degree of the expulsion defects to that of *hp684* allele (*hp684* 7.55 ± 0.34/10 min; *hp821* 6.8 ± 0.36/10 min; *hp821hp833* 7.6 ± 0.47/10 min; *hp865* 8 ± 0.33/10 min). To briefly describe the change, we used “Exp/Cycle” in the following quantifications (WT 0.98 ± 0.01; *hp684* 0.55 ± 0.02; *hp821* 0.49 ± 0.03; *hp821hp833* 0.53 ± 0.03; *hp865* 0.57 ± 0.03) (Wang *et al*., 2013). All alleles also selectively regulated the expulsion frequency without affecting the pBoc and aBoc steps (**Fig 1D and EV1C, D**); the UBR-1 genomic fragment rescued expulsion frequency in all alleles (**Fig 1D and EV1C, D**).

These results demonstrate that the ∼50% reduction of expulsion represents the effect of a full functional loss of the UBR-1 protein. It’s significant to note that we found that the expression of the human ortholog of UBR1 restored the reduced expulsion in *ubr-1* loss-of-function mutants (**Fig 1D**), indicating functional conservation. We analyzed the *hp684* allele in most follow-up experiments.

### *ubr-1* promotes the expulsion motor step through the GABAergic neuron that contracts enteric muscles

UBR-1 expression has been observed consistently in neurons and muscles (Chitturi *et al*., 2018; Hwang *et al*, 2011; Kwon *et al*, 2001). To pinpoint the tissue requirement for expulsion, we used tissue-specific promoters to restore UBR-1 expression in the *ubr-1* mutants.

The expulsion step is carried out by contraction of enteric muscles, driven by two classes of excitatory GABAergic neurons DVB and AVL (**Fig 1A**) (Branicky & Hekimi, 2006; McLntire *et al*, 1993b; Thomas, 1990). We found that a pan-neuronal, but not a pan-muscle expression of UBR-1, fully restored the frequency of expulsion (**Fig 1E and F**). Restoration of UBR-1 in the DVB, AVL and D-class GABAergic neurons (D-MNs, P*unc-47*, 0.97 ± 0.01 exp/cycle) or in the DVB, AVL and RIS neurons (P*lim-6 (int3)*, 0.94 ± 0.02 exp/cycle) (Hobert *et al*, 1999) also fully restored the defecation frequency of *ubr-1* mutants (**Fig 1E and F, and EV2A**). When we restored UBR-1 expression exclusively in the DVB neuron (P*flp-10*, 0.92 ± 0.03 exp/cycle) (Choi *et al*, 2021), the expulsion defect of *ubr-1* mutant was also rescued. By contrast, expression of UBR-1 in RIS neuron (P*flp-11*, 0.61 ± 0.01 exp/cycle) (Turek *et al*, 2016), the expulsion defect was not rescued (**Fig 1E and F, and EV2B**), similar to restoring UBR-1 expression in either D-class neurons (P*unc-25s*, 0.57 ± 0.06 exp/cycle) (Jin *et al*, 1999), cholinergic neurons (P*acr-2*, 0.54 ± 0.04 exp/cycle) or interneurons (P*glr-1*, 0.49 ± 0.05 exp/cycle). Thus, the *ubr-1* mutant’s expulsion defect likely reflects functional defects in DVB and AVL, the GABAergic neurons that mediate the enteric muscle contraction. Consistently, both the translational reporter (**Fig EV2C**) and the transcriptional reporter (**Fig 1G and EV2D**) for the *ubr-1* gene exhibited strong and consistent expression in DVB and AVL neurons.

Collectively, UBR-1 promotes the expulsion motor step through the GABAergic neurons that contract enteric muscles for defecation.

### UBR-1 is required for DVB neuron’s oscillated calcium activity

Previous studies revealed that DVB neuron exhibits periodic Ca^2+^ activities that are tightly correlated with the expulsion behavior (Jiang *et al*, 2022; Wang *et al*., 2013). We asked whether the reduced expulsion of *ubr-1* mutants is associated with changes in DVB activity.

Using a genetic calcium sensor GCaMP6s (green) for DVB with a calcium insensitive wCherry (red) as reference (**Fig 2A and EV3A**), we examined the Ca^2+^ activity in DVB neuron. We observed robust oscillation Ca^2+^ transients in loosely glued wild type animals (**Fig EV3B and C, Methods**), similar to that in free-moving animals (Wang *et al*., 2013). These Ca^2+^ transients exhibited same frequency and amplitude in either dual-channel (GCaMP6s/wCherry, 5.4 ± 0.65 Hz/180s) or single-channel (GCaMP6s, 4.63 ± 0.17 Hz/180s) Ca^2+^ imaging (**Fig 2B and EV3D**). For simplicity, subsequent experiments were performed with single-channel. We found that each epoch of DVB’s Ca^2+^ transient was accompanied by an execution of the expulsion step (**Movies EV1**). Furthermore, the expulsion action and DVB Ca^2+^ oscillation were simultaneously abolished in glued *nlp-40* mutant (data not shown), an instructive neuropeptide from the intestine that delivers temporal information to the DVB neuron (Wang *et al*., 2013), suggesting that the DVB Ca^2+^ dynamics are dependent on NLP-40 initiated pacemaker signal.

**Figure 2.**
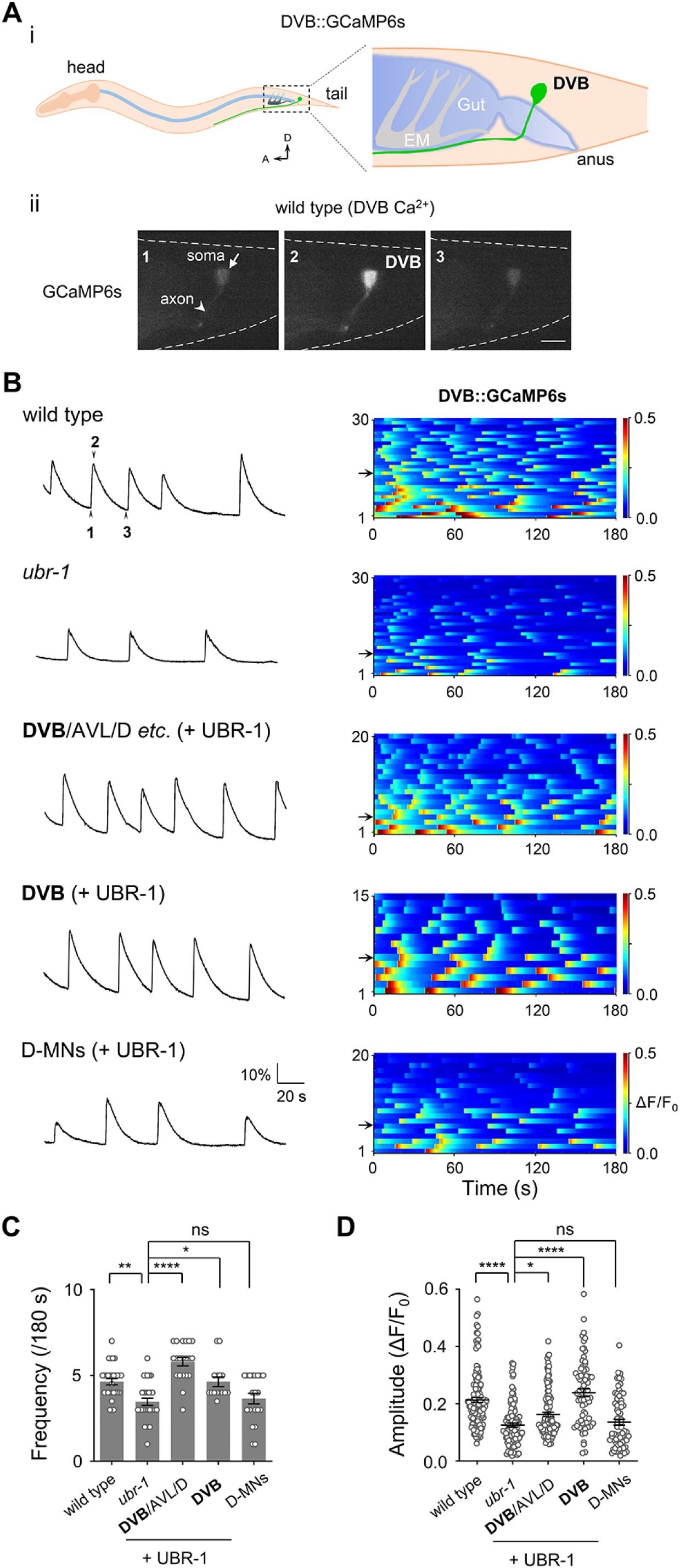
*ubr-1* regulates DVB Ca^2+^ activity. (A) Schematic diagram of DVB neuron calcium imaging. (i) Schematic DVB::GCaMP6s showing cell body and axon in the tail. (ii) Three sequential real-time fluorescence snapshots of a single Ca^2+^ transient (B) in wild type animal. Scale bar, 10 µm. (B) *Left*, representative DVB rhythmic Ca^2+^ transient traces in different genotypes. *Right*, color maps summaries the DVB Ca^2+^ activity. Each horizontal line corresponds to one animal of the respective genotypes. ΔF/F_0_ = (F-F_0_)/F_0_ was calculated. (C, D) Quantification of the frequency and peak amplitude of DVB Ca^2+^ transients. *ubr-1* mutants exhibited significant reduced frequency and amplitude, which were capable of rescued by restoring UBR-1 expression in DVB and AVL neurons, but not in D-motor neurons. \*\**p* < 0.01; \*\*\**p* < 0.001; \*\*\*\**p* < 0.0001; one-way ANOVA test. n = 15-20 animals. Error bars, SEM.

In *ubr-1* mutants, DVB neuron exhibited reduced frequency (3.46 ± 0.21 Hz/180s) (**Fig 2B and C**), and the peak amplitude (**Fig 2B and D**) of the Ca^2+^ oscillatory transients. Morphological development of the DVB neuron was similar between the wild type and *ubr-1* animals (**Fig EV4A**). The loss of UBR-1 therefore compromises DVB neuronal activity.

Consistent with the behavioral observations, a restored expression of UBR-1 in DVB, AVL and D-MNs (P*unc-47*) or in DVB (P*flp-10*), but not in D-MNs (P*unc-25s*), fully restored the frequency and peak amplitude of DVB’s Ca^2+^ oscillatory transients (**Fig 2B-D**). These results support the notion that that UBR-1 regulates DVB activity cell-autonomously.

Although AVL’s activity was not examined directly, the known gap junctional connection between AVL and DVB supports the notion that AVL activity should be co-regulated by UBR-1 (Bhattacharya *et al*, 2019; Choi *et al*., 2021).

### *ubr-1* mutant’s expulsion defect is not due to insufficient GABAergic signaling

DVB is a GABAergic motor neuron (McLntire *et al*., 1993b). It releases GABA, which activates EXP-1, a GABA-gated cation channel expressed by the enteric muscles to activate muscle contraction (Beg & Jorgensen, 2003). The simplest implication of the reduced expulsion and DVB Ca^2+^ activity in *ubr-1* mutants is a defective GABAergic signaling the system, due to either reduced GABA release and/or GABA receptor EXP-1.

To test these possibilities, we examined whether the expulsion frequency of *ubr-1* mutants could be restored by exogenous GABA application. Consistent with previous report (McLntire *et al*, 1993a), addition of GABA to the culture media significantly increased the expulsion frequency of *unc-25*, a mutant that lacks the enzyme to convert glutamate to GABA (**Fig EV4B**) (Jin *et al*., 1999). However, the expulsion defect of the *ubr-1* mutant was not improved by exogenous GABA application (**Fig EV4B**).

We further compared the endogenous GABA level of synchronized wild-type and *ubr-1* young adults by high-performance liquid chromatography (HPLC). To our surprise, but consistent with the observation of exogenous GABA application, the GABA level not only did not exhibit a decrease, but instead showed a moderate increase in *ubr-1* mutants (**Fig EV4C**). An increase of GABA level is also consistent with the metabolism consequence of an elevated glutamate in *ubr-1* mutants (Chitturi *et al*., 2018). Lastly, the localization and expression density of the postsynaptic GABA receptor EXP-1 showed no obvious change *in ubr-1* mutants (**Fig EV4D**).

Thus, the reduced defecation frequency or DVB neuronal activity of the *ubr-1* mutant could not be explained by insufficient excitatory GABA signaling.

### Preventing glutamate release in *ubr-1* mutants restores the defecation frequency

If it is not through GABA signaling, how might UBR-1 affects DVB activity? Glutamate, GABA’s sole precursor (Miller *et al*, 1978) is elevated in *ubr-1* mutants (Chitturi *et al*., 2018). We reason that an increased glutamate signaling might contribute to *ubr-1*’s defecation defects.

We examined the effect of blocking glutamate release in the *ubr-1* mutant background by removing EAT-4, the vesicular glutamate transport (Lee *et al*, 1999). Remarkably, removal of EAT-4 robustly restored *ubr-1*’s expulsion frequency (*ubr-1; eat-4*, 12 ± 0.30/10 min) (**Fig 3A and B**). Conversely, overexpression of EAT-4 in *ubr-1* mutant (P*unc-47*::EAT-4) worsen the defecation defect (**Fig EV4E**). These results suggest that glutamate signaling might directly regulate expulsion. This was unexpected because previous studies did not reveal synaptic connections from glutamatergic neurons to enteric muscles. We found that a transcriptional reporter of *eat-4* showed moderate but consistent expression in DVB and AVL (**Fig 3C**). This suggests the possibility of dual transmission of glutamate and GABA, as observed in other systems (Root *et al*., 2018), from these enteric muscle-innervating, excitatory GABAergic neurons.

**Figure 3.**
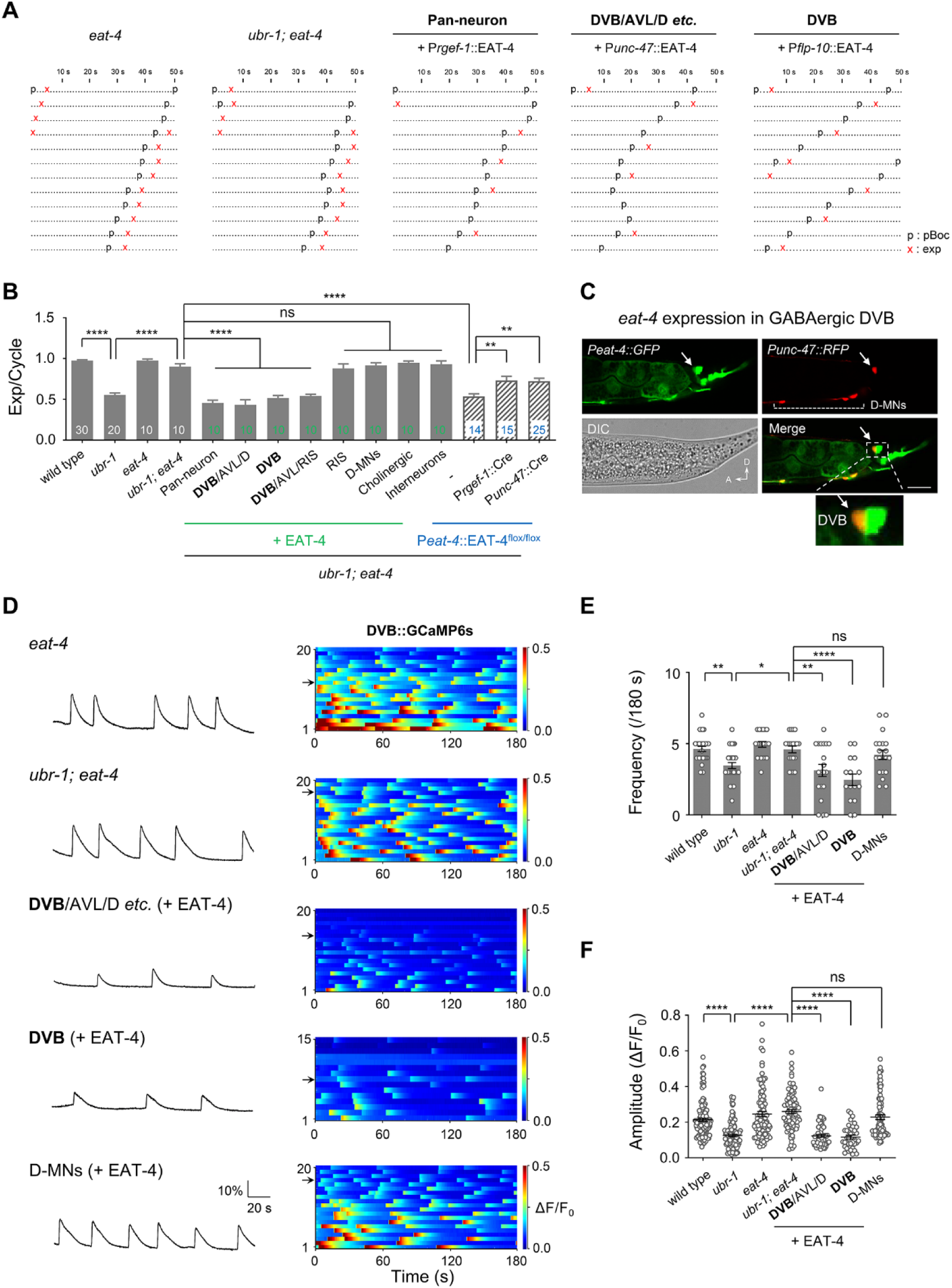
*eat-4* suppresses *ubr-1* expulsion defect. (A) Representative ethograms of consecutive 10 min defecation cycles in indicated genotypes. Each dot represents 1 s. ‘‘p’’ and ‘‘x’’ stand for pBoc and exp, respectively. (B) Quantification of the frequency of expulsion in all genotypes. Loss of function mutants in *eat-4* suppress *ubr-1* expulsion defect. DVB specific restoring of EAT-4 reverted expulsion frequency back to *ubr-1*. Cre-LoxP knock-down of *eat-4* expression in DVB suppressed the rescue of *ubr-1; eat-4* mutants by P*eat-4*::EAT-4^flox/flox^, which has *loxp* flanked (floxed) allele of *eat-4* minigene. \*\**p* < 0.01; \*\*\*\**p* < 0.0001; one-way ANOVA test. (C) P*eat-4*::GFP expression in DVB neuron. White arrow denotes the DVB soma, and D-motor neurons are labeled. Scale bar, 20 µm. (D) Representative DVB Ca^2+^ transient traces and color-maps of indicated genotypes. (E, F) Quantification of the frequency and peak amplitude of DVB Ca^2+^ transients. *eat-4* suppresses the DVB Ca^2+^ frequency and amplitude of *ubr-1* mutants. \**p* < 0.05; \*\**p* < 0.01; \*\*\*\**p* < 0.0001; one-way ANOVA test. n = 10-30 animals. Error bars, SEM.

Further supporting the possibility that increased glutamate signaling in the defecation circuit might be the cause of *ubr-1*’s reduced defecation, restoring EAT-4 expression in DVB, AVL and D-class GABAergic neurons (P*unc-47*, 0.90 ± 0.03 exp/cycle) or in DVB, AVL and RIS neurons (P*lim-6 (int3)*, 0.54 ± 0.02 exp/cycle) or in single DVB neuron (P*flp-10*, 0.51 ± 0.03 exp/cycle) in *ubr-1; eat-4* mutants reverted the expulsion frequency back to that of *ubr-1* mutants (0.55± 0.02 exp/cycle) (**Fig 3B**). Reversion was not observed when we drove EAT-4 expression in RIS neuron (P*flp-11*, 0.88 ± 0.05 exp/cycle), D-class neurons (P*unc-25s*, 0.92 ± 0.03 exp/cycle), cholinergic neurons (P*acr-2s*, 0.94 ± 0.02 exp/cycle), or premotor interneurons (P*glr-1*, 0.93 ± 0.04 exp/cycle). Lastly, when we knock-down *eat-4* expression in DVB and AVL (Methods) in *ubr-1* mutants, expulsion frequency was significantly increased (0.72 ± 0.04 exp/cycle) (**Fig 3B**). These results establish a direct role of vesicular glutamate transporter EAT-4 in the defecation circuit.

Recapitulating the genetic interactions at the behavioral level, removing *eat-4* in *ubr-1* mutant background also led to increased DVB calcium activity (**Fig 3D-F**). Restoring EAT-4 in DVB, AVL and D, or in DVB alone, but not in D in *ubr-1; eat-4* mutants also reverted DVB’s calcium activity to that of *ubr-1* mutants (**Fig 3D-F**). Thus, the neuronal activity could also be compensated by the removal of VGLUT in these GABAergic motor neurons. Consistent with behavioral defect, overexpression of EAT-4 in *ubr-1* mutant (P*unc-47*::EAT-4) further decreased the DVB calcium activity (**Fig EV4F and G**). We observed no significant difference in *eat-4* transcription and EAT-4 abundance level between *ubr-1* and and wild-type animals (**Fig EV4H-J**), suggesting that *eat-4* is not regulated by UBR-1.

Neither the expulsion frequency or DVB activity was changed between *eat-4* loss of function mutants and wild-type animals (**Fig 3B, E and F**). The pBoc frequency in *eat-4*, or *eat-4; ubr-1* mutants was not altered, indicating that the DMP cycle interval are normal in these mutants (**Fig EV4K**). This implicates a low basal glutamatergic signaling at the defecation circuit, which might play a modulatory role.

### Inhibitory neuronal glutamatergic signaling contributes to *ubr-1* mutant’s reduced expulsion

A glutamatergic signaling from DVB should activate glutamate receptors of the defecation circuit. *C. elegans* genome encodes four classes of glutamate receptors, including the excitatory classes (the ionotropic AMPA and NMDA-type), the metabotropic classes (mGluRs), and the inhibitory classes (the glutamate-gated chloride channels GluCls) (Brockie & Maricq, 2006; Dillon *et al*, 2006).

To address which glutamate receptors underlies glutamatergic signaling in the defecation circuit, we began with examining their expression patterns. Excitatory glutamate receptors have been reported to be expressed in the nerve system, but none was reported in the expulsion circuit (**Table S1**) (Brockie *et al*, 2001; Greer *et al*, 2008; Jeong & Paik, 2017; Katz *et al*, 2019). The expression pattern of inhibitory GluCls was not as comprehensively described (Holden-Dye & Walker, 2014). We generated transcriptional reporters of all six GluCls and found that one, the *glc-3* reporter exhibited strong expression in DVB (**Fig 4A and EV5**).

**Figure 4.**
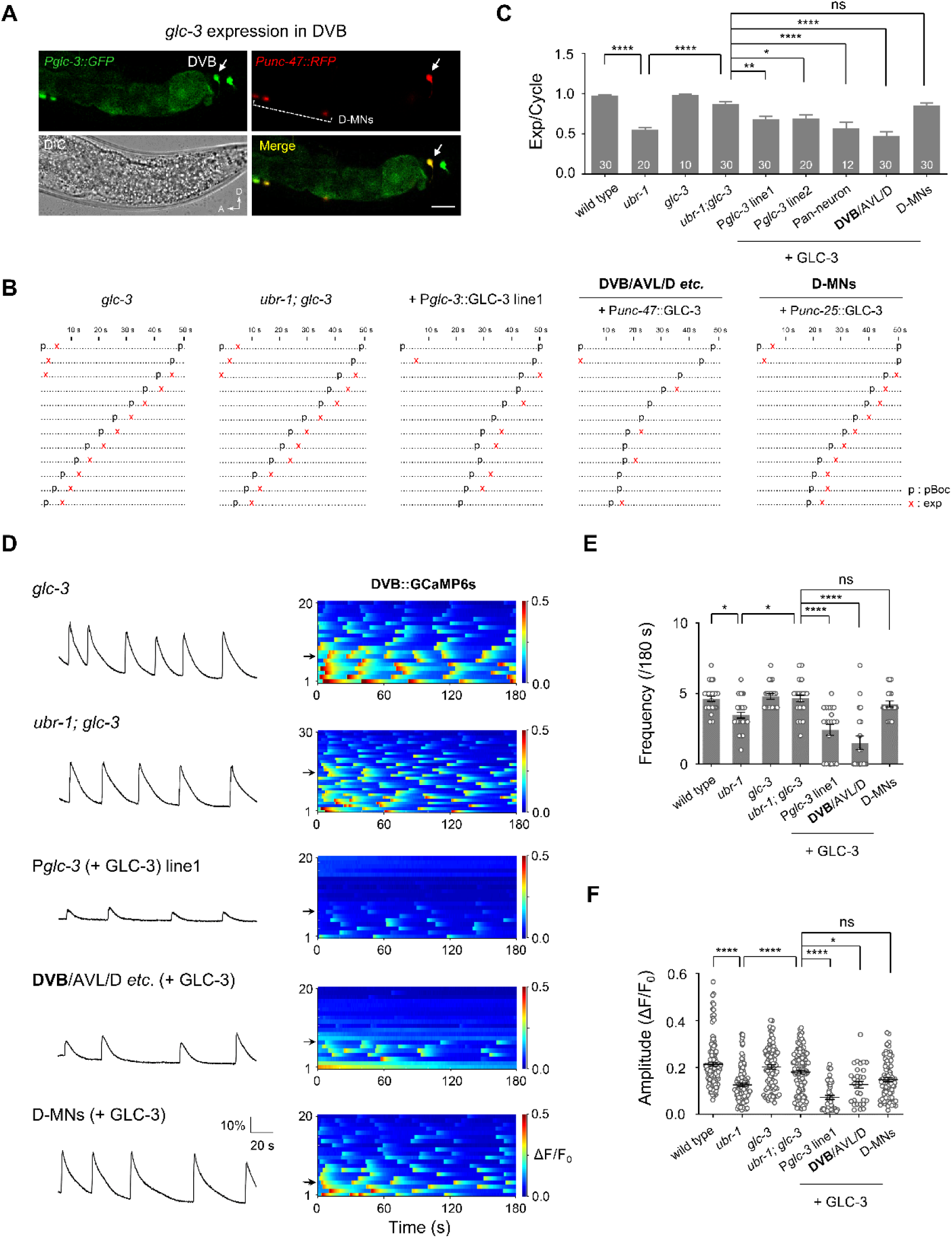
Glutamate-gated chloride channel GLC-3 regulate UBR-1-mediated expulsion. (A) Expression of P*glc-3*::GFP in DVB neuron. Scale bar, 20 µm. (B) Representative ethograms of 10 min defecation cycles of indicated genotypes. Each dot represents 1 s. ‘‘p’’ and ‘‘x’’ stand for pBoc and exp, respectively. (C) Quantification of the frequency of expulsions in different genotypes. Loss of function mutations in *glc-3* suppress *ubr-1* expulsion defect. Restoring GLC-3 in DVB and AVL neurons reverted the suppression of *glc-3* to *ubr-1*. \*\**p* < 0.01; \*\*\*\**p* < 0.0001; one-way ANOVA test. (D) Representative DVB Ca^2+^ transient traces (*left*) and all Ca^2+^ activity color-maps (*right*) of indicated genotypes. (E, F) Quantification of the frequency and peak amplitude of the Ca^2+^ transients. *glc-3* cell-autonomously suppresses the DVB Ca^2+^ frequency and amplitude of *ubr-1* mutants. \**p* < 0.05; \*\**p* < 0.01; \*\*\*\**p* < 0.0001; one-way ANOVA test. n = 10-30 animals. Error bars, SEM.

Co-presence of the glutamate release machinery (EAT-4) and the inhibitory glutamate receptor (GLC-3) in DVB implies an auto-inhibitory regulation. To test this possibility, we examined the effect of removing GLC-3. In *ubr-1; glc-3* mutants, the expulsion frequency (0.87 ± 0.03 exp/cycle), as the case for *ubr-1; eat-4*, was significantly rescued (**Fig 4B and C**). Furthermore, restoring GLC-3 expression, either pan-neuronally or in DVB and AVL, but not in D, reverted the expulsion frequency of *ubr-1; glc-3* to that of *ubr-1* mutants (**Fig 4B and C**). DVB’s calcium activity in these mutants fully recapitulated the behavioral effects of the corresponding genotypes (**Fig 4D-F**).

Similar to *eat-4* mutants, we did not find significant differences in pBoc frequency, expulsion frequency or DVB activity between *glc-3* mutants and wild-type animals (**Fig 4C and EV4K**). These results implicate that glutamate signaling from DVB negatively regulates its activity, an effect that is likely amplified by elevated glutamate level in *ubr-1* mutants.

### Inhibitory glutamatergic signaling from DVB contributes to *ubr-1*’s expulsion defect

We further noted that the transcription reporters for *glc-2* and *glc-4* showed strong expression in muscles of the defecation circuit, including the enteric and anal depressor muscles (**Fig 5A and B, and EV5**). These muscles are responsible for expulsion (Branicky & Hekimi, 2006). We examined whether they also contribute to the expulsion defect of *ubr-1* mutants.

**Figure 5.**
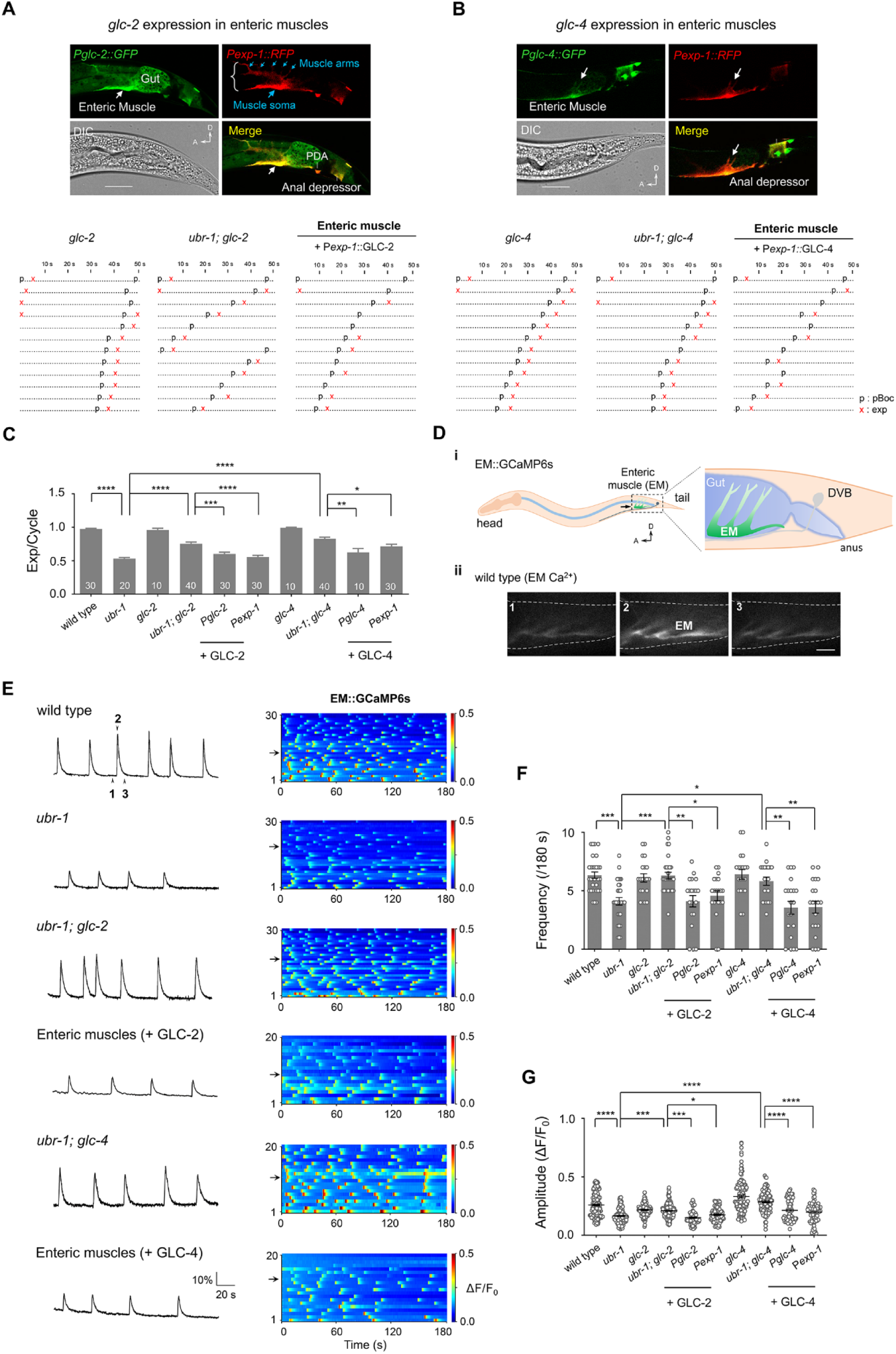
Intestinal Muscular GLC-2/4 are required for UBR-1-mediated expulsion. (A, B) *Upper*, expression patterns of P*glc-2*::GFP (*left*) and P*glc-4*::GFP (*right*) show enteric muscles (EM) localizations. The muscle soma and muscle arms were denoted by blue arrows. Scale bar, 20 µm. *Bottom*, representative ethograms of consecutive 10 min defecation cycles of indicated genotypes. Each dot represents 1 s. ‘‘p’’ and ‘‘x’’ stand for pBoc and exp, respectively. (C) Quantification of the expulsion frequency in different genotypes. Loss of function mutations in *glc-2* or *glc-4* suppress *ubr-1* expulsion defect. Restoring GLC-2 or GLC-4 in enteric muscles reverted the expulsion suppression of *glc-2/4* to *ubr-1*. \**p* < 0.05; \*\**p* < 0.01; \*\*\**p* < 0.001; \*\*\*\**p* < 0.0001; one-way ANOVA test. (D) (i) Schematic EM::GCaMP6s showing cell soma (EM) and muscle arms around the posterior intestine (Gut) in the tail. (ii) Three sequential real-time fluorescence snapshots of a single EM Ca^2+^ transient in wild type (E). Scale bar, 20 µm. (E) *Left*, representative EM Ca^2+^ traces in different genotypes. *Right*, color maps represent all EM Ca^2+^ activity. Each horizontal line corresponds to one animal of the respective genotypes. ΔF/F_0_ = (F-F_0_)/F_0_ was calculated. *ubr-1* mutants exhibited reduced EM Ca^2+^ activity, which was suppressed by *glc-2/4*. (F, G) Quantification of the frequency and peak amplitude of EM::Ca^2+^ transients. Restoring GLC-2 and GLC-4 rescued frequency and amplitude of intestinal muscle activation in *ubr-1 glc-2* and *ubr-1; glc-4* mutants, respectively. \**p* < 0.05; \*\**p* < 0.01; \*\*\**p* < 0.001; \*\*\*\**p* < 0.0001; one-way ANOVA test. n = 10-40 animals. Error bars, SEM.

The functional loss of either *glc-2* or *glc-4* in *ubr-1* mutants led to an increase of their expulsion frequency (*ubr-1; glc-2* 0.75 ± 0.03 exp/cycle; *ubr-1; glc-4* 0.83 ± 0.02 exp/cycle) (**Fig 5A-C**). Restoring GLC-2 or GLC-4 to the enteric muscles in *ubr-1 glc-2* or *ubr-1; glc-4* mutants reverted their expulsion frequencies to that of *ubr-1* mutants (**Fig 4B and C**). These results implicate an inhibitory glutamatergic signaling, from DVB to the enteric muscles contributes to the expulsion defect of *ubr-1* mutants.

To directly examine the role of GLC-2 and GLC-4 on expulsion, we examined the enteric muscle’s calcium activity (**Fig 5D**). In wild-type animals, as expected, enteric muscles exhibited robust, oscillating calcium signals tightly associated with expulsion (**Fig 5D and E, and Movies EV2**). In *ubr-1* mutants, enteric muscles exhibited a significant reduction in the frequency and peak amplitude of calcium signals, reminiscent of the characteristics of reduced DVB activity (**Figure 5E**), and the decreased enteric muscle activity associated with reduced expulsion frequency in *ubr-1* mutants. Enteric muscles showed normal shape and comparable basal GCaMP intensity between wild type and *ubr-1* animals (**Fig EV4A**).

Fully corroborating the effect on behaviors, the enteric muscle’s calcium signals in *ubr-1; glc-2* mutants exhibited an increase compared to *ubr-1* mutants (**Fig 5E-G**), and effect reverted when we restored the expression of GLC-2 in enteric muscles (**Fig 5E-G**). We observed similar genetic interactions between *ubr-1* and *glc-4* (**Fig 5E-G**). These results implicate that GLC-2 and GLC-4 may form a heteromeric GluCl receptor to negatively regulate the activity of enteric muscles.

Similar to *eat-4* and *glc-3* mutants, we did not find changes in pBoc frequency in *glc-2*, *ubr-1; glc-2* or *glc-4* and *ubr-1; glc-4* compare to wild type (**Fig EV4K**), although the expulsion frequency was partially restored when compare *ubr-1; glc-2* to *glc-2*, or *ubr-1; glc-4* to *glc-4* animals (**Fig 4C**). These results indicate that an elevated inhibitory glutamate signaling between the DVB motor neuron and enteric muscles, in response to increased glutamate release in *ubr-1* mutants, contributes to the mutant animal’s reduced expulsion.

### Glutamate perfusion potently inhibits the defecation circuit

A global elevation of glutamate level was observed in *ubr-1* mutants (Chitturi *et al*., 2018). Whether this leads to an increase of the functional neurotransmitter in the defecation circuit has not been demonstrated directly.

We first performed glutamate perfusion to mimic the effect of excessive glutamate. When animals were exposed to glutamate, DVB’s calcium activity was significantly reduced (**Fig 6A**). Quantitatively, at 5.9 mM, exogenous glutamate exposure reduced the frequency (**Fig 6B**) and peak amplitude (**Fig 6C**) of calcium signals from DVB. Importantly, the same glutamate treatment did not lead to inhibition in *glc-3* mutants (**Fig 6B and C**), consistent with the notion that exogenous glutamate inhibits DVB activity through GLC-3. Similarly, such glutamate exposure reduced the activity of enteric muscles, with a modestly reduced frequency (**Fig 6D and E**) and significantly reduced amplitude (**Fig 6F**) of DVB’s calcium transient. Glutamate-induced inhibition of enteric muscle was not observed in *glc-2*; *glc-4* double mutants (**Fig 6E and F**), which also exhibit significant suppression of *ubr-1* (data not shown). Together, they support the notion that increased glutamate could lead to increased inhibitory glutamate signaling in the defection circuit.

**Figure 6.**
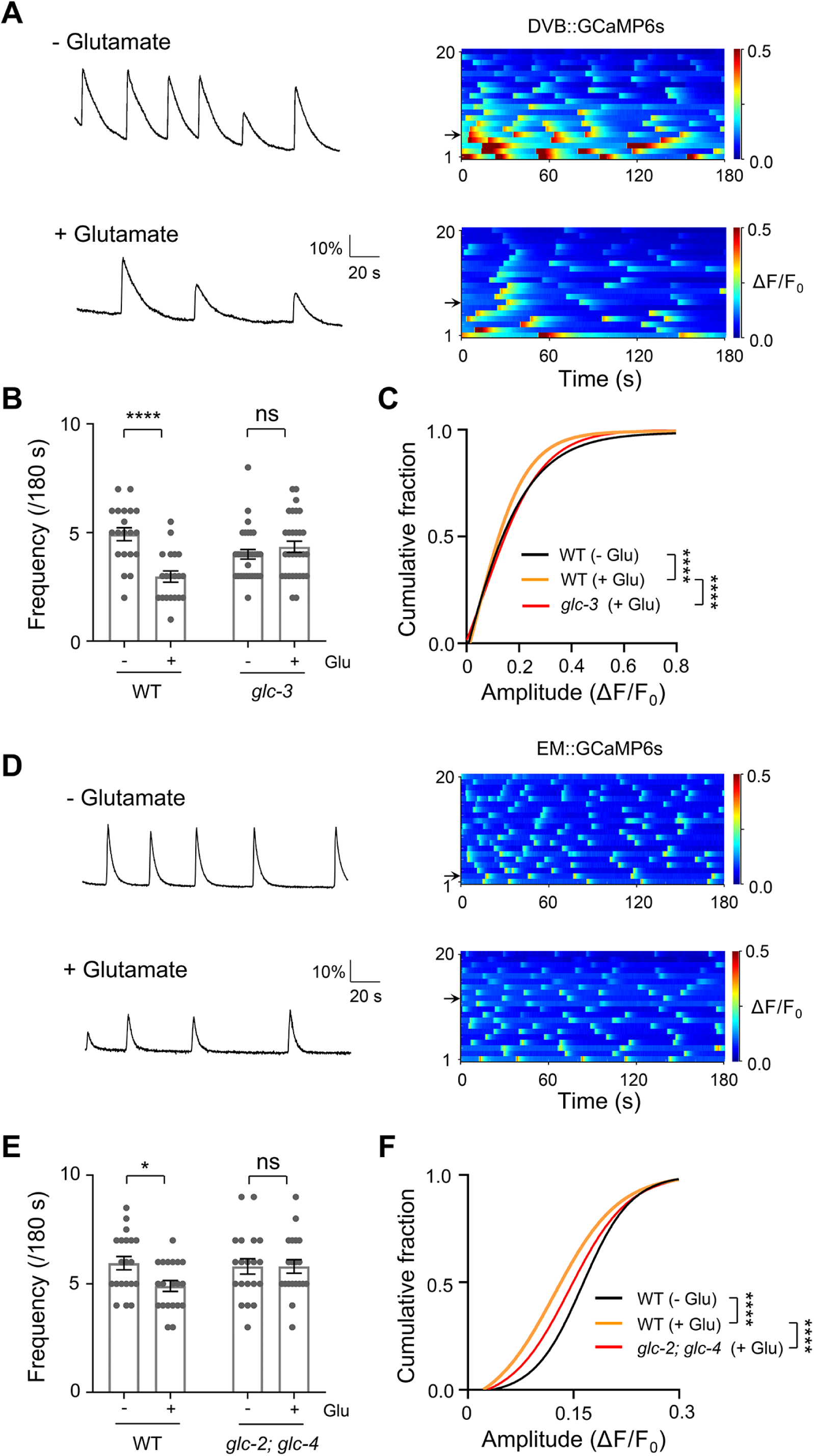
Glutamate inhibits DVB neuron and enteric muscles through the activation of GLC-3 and GLC-2/GLC-4. (A) Representative DVB Ca^2+^ traces (*left*) and Ca^2+^ activity color-maps (*right*) in wild-type animals without and with glutamate. ΔF/F_0_ = (F-F_0_)/F_0_ was calculated. Glutamate significantly reduced the DVB Ca^2+^ activity. (B) Quantification of the frequency of DVB Ca^2+^ transients. Wild-type animals exhibited significant reduced frequency, while *glc-3* mutant did not change after glutamate application. \*\*\*\**p* < 0.0001; two-way ANOVA test. (C) Distribution of the amplitude of DVB Ca^2+^ transients in wild-type and *glc-3* mutant, with and without glutamate perfusion. Glutamate leads to a reduction of amplitude in wild-type animals, but not in *glc-3* mutant. \*\*\*\**p* < 0.0001; Kolmogorov-Smirnov test. (D) *Left,* representative enteric muscles (EM) Ca^2+^ transient traces in wild-type and glutamate perfusion animals. *Right*, color maps represent EM Ca^2+^ activity. Glutamate reduced EM Ca^2+^ activity. (E) Quantification of the frequency of EM Ca^2+^ transients. Wild-type animals exhibited significant reduced frequency, while *glc-2; glc-4* mutant did not change after glutamate application. \**p* < 0.05; two-way ANOVA test. (F) Distribution of the amplitude of EM Ca^2+^ transients in wild-type and *glc-2; glc-4* mutant, with and without glutamate perfusion. Glutamate leads to a reduction of amplitude in wild-type animals, but not in *glc-2; glc-4* mutant. \*\*\*\**p* < 0.0001; Kolmogorov-Smirnov test. n = 20-30 animals. Error bars, SEM.

To further determine that glutamate may act as a signaling molecule at the defection circuit, we puffed glutamate onto the DVB neuron in a dissected *C. elegans* preparation (**Fig 7A and Methods**). In wild-type dissected preparation, DVB exhibited robust oscillatory calcium signals in a bath solution with high chloride (167 mM) (**Methods**). 1 mM glutamate perfusion led to an instantaneous and potent attenuation of DVB’s calcium signals (**Fig 7B and C**). This attenuation was reversed with washout of a glutamate-free solution, indicating that glutamate directly inhibits DVB. Importantly, the inhibitory effect of glutamate was abolished in low chloride (17 mM) (**Fig 7B and C**), indicating glutamate activation of a chloride-dependent inhibitory conductance on DVB. In *glc-3* mutants, DVB’s calcium signals was not affected by glutamate puff (**Fig 7B and C**). These results strongly support the presence of an auto-inhibitory glutamate signaling at DVB through GLC-3.

**Figure 7.**
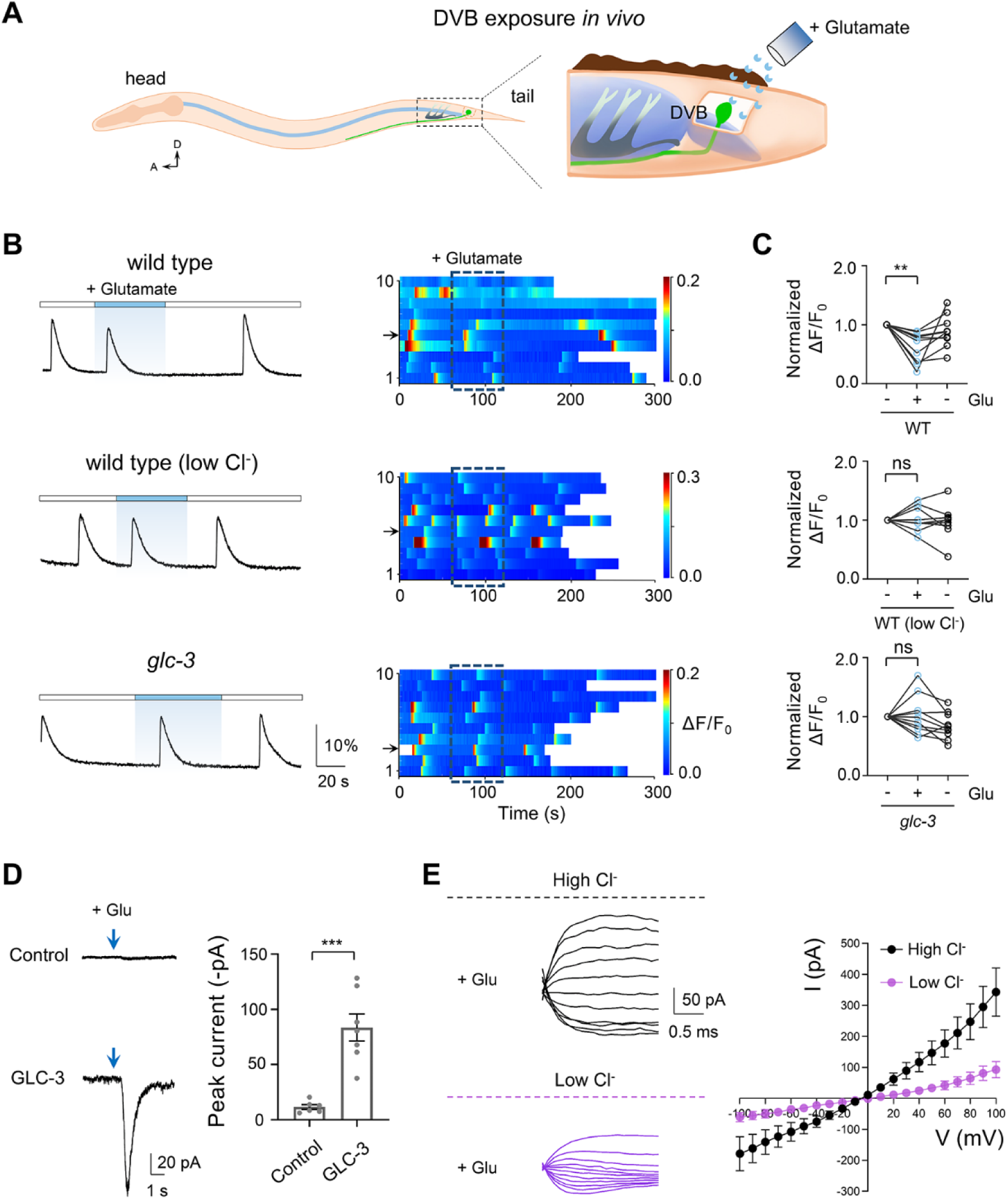
Glutamate directly inhibits dissected DVB neuron. (A) Schematic diagram of dissected DVB neuron *in vivo*. (B) Robust rhythmic Ca^2+^ transients were recorded from the dissected DVB neuron. Representative DVB Ca^2+^ transient traces (*left*) and Ca^2+^ activity color-maps (*right*) of wild-type and *glc-3* mutant. (C) Extracellular fluid with glutamate (1 mM) significantly reduces the peak amplitude of DVB Ca^2+^ activity, which is lost in low Cl^−^ bath solution and *glc-3* mutant. \*\**p* < 0.01; one-way ANOVA test. (D) Responses to glutamate of HEK293T cells expressing empty vector and GLC-3. Glutamate evokes inward currents in cells expressing *C. elegans* GLC-3. \*\*\**p* < 0.001; two-tailed unpaired *t*-test. (E) Representative step currents were recorded by whole-cell voltage-clamp from HEK293T cells with expression of GLC-3. Cells were depolarized from −80mV to +80mV with an increment of +40 mV. (F) *Left*, representative step currents were recorded with high Cl^−^ (150.6 mM) and low Cl^−^ (10.6 mM) concentration in bath solution. *Right*, Cells were depolarized from −100mV to +100mV with an increment of +20 mV. I-V curve with the expression of GLC-3. n = 10 animals in each group. Error bars, SEM.

GLC-3 forms a glutamate-gated chloride-dependent conductance, which we further reconstituted in the HEK293T cells. In these cells, 1 mM glutamate activated a GLC-3-dependent inward current at −60 mV in the bath solution with high chloride (150.6 mM) (**Fig 7D**). This Glutamate-activated and GLC-3-dependent conductance was significantly decreased in low extracellular chloride (10.6 mM) (**Fig 7E**).

These results demonstrate that the glutamate acts as an inhibitory signaling to negatively regulate the defecation circuit activity.

## Discussion

We reveal that the UBR-1 E3 ligase regulates the relative excitatory/inhibitory signaling strength at the defecation motor circuit. This is an excitatory GABAergic defecation circuit that also has a low level of inhibitory glutamatergic signaling. This signaling consists of the GABAergic motor neuron DVB, which also releases glutamate, with an auto-inhibitory signaling of DVB and descending inhibition of enteric muscles by glutamate-gated chloride channels. In *ubr-1* mutants, elevated glutamate leads to increased inhibitory signaling thus reduced defecation circuit activity and expulsion.

### A functional model for UBR-1-mediated regulation of E/I balance at the defecation circuit

We propose a functional model of the defecation circuit and UBR-1’s effect (**Synopsis**). In these GABAergic neurons, there is dual - excitatory GABAergic and inhibitory glutamatergic - signaling, where the GABAergic signaling plays the dominant role. UBR-1 maintains this signaling imbalance. When UBR-1 is dysfunctional, glutamate level is increased, which leads to elevated glutamate upload and release, consequently an increased inhibitory glutamatergic signaling. This increase impedes the functional output of the excitatory GABAergic signaling, reducing the functional output, the frequency of expulsion.

Mechanistically, GABAergic signaling excites the defecation circuit through DVB-mediated GABA release and EXP-1-mediated activation of the enteric muscles. On other hand, glutamatergic signaling inhibits the defecation circuit through GLC-3-mediated DVB auto-inhibition and GLC-2/4-mediated enteric muscle inhibition. By gating the glutamate level, UBR-1 regulates the relative strength of excitatory GABAergic and inhibitory glutamatergic signaling, affecting DVB activity and final output of the defecation motor program, expulsion.

### The potential role of dual signaling: a robust motor program with flexibility

The defecation motor program is highly stereotypic, with robust short-term rhythm (Thomas, 1990), and the underlying signaling pathway have been well characterized (Branicky & Hekimi, 2006; Choi *et al*., 2021; Dal Santo *et al*, 1999; Jiang *et al*., 2022; Mahoney *et al*, 2008; McLntire *et al*., 1993a; McLntire *et al*., 1993b; Nehrke *et al*, 2008; Teramoto & Iwasaki, 2006; Wang *et al*., 2013). This motor program can be reset by food content and mechanical stimulation (Liu & Thomas, 1994; Thomas, 1990), hence this robust motor circuit also allows modulation. Our study reveals one circuit mechanism for modulation.

DVB drives the excitatory GABAergic signaling (McLntire *et al*., 1993b) that drives the periodic enteric muscle contraction (Beg & Jorgensen, 2003). Its co-release of glutamate, which activates inhibitory signaling, serves a modulatory role to fine-tune the rhythm output. Unlike the sensorimotor circuit, which offers flexibility by coordinating the excitatory and inhibitory signaling through different neurotransmitters from discrete neurons (Zhen & Samuel, 2015) or different receptors to the same neurotransmitter (Sato *et al*, 2021), DVB utilizes a dual-neurotransmitter system of opposite signs, glutamate and GABA, to regulate this stereotyped motor function. This allows the compact defecation motor circuit to make adaptive changes. The dual signaling from DVB has a strong dominance of excitatory GABAergic signaling over low level inhibitory glutamatergic signaling. This GABAergic/glutamatergic signaling ratio is likely critical for the robustness of the defecation motor program to exhibit flexibility.

### Co-release of GABA and glutamate

In the mammalian brain, neurons have been shown to produce and release multiple neurotransmitters. Inhibitory GABA and excitatory glutamate have been reported to co-exist and being co-released from the same synapses in the central nervous system (Barker *et al*, 2017; Kim *et al*, 2022; Root *et al*, 2014; Root *et al*., 2018; Shabel *et al*., 2014). The GABA/glutamate co-release was altered in animal models of depression and addiction (Meye *et al*, 2016; Shabel *et al*., 2014), highlighting its functional and behavioral relevance.

GABA can also be an excitatory neurotransmitter in the immature brain (Ben-Ari, 2002; Owens & Kriegstein, 2002). Na^+^-dependent GABA depolarizations have also been documented in the adult nervous system, such as the stomatogastric ganglion of the crab and the accessory olfactory bulb of the rat (Chavas & Marty, 2003; Goldmakher & Moss, 2000; Gulledge & Stuart, 2003; Lu & Trussell, 2001; Swensen *et al*, 2000). The inhibitory effects of glutamate effect through mGluRs have also been reported (Pinheiro & Mulle, 2008).

Our study offers a new example of the co-release of excitatory GABA and inhibitory glutamate. This new functional mode occurs not only by the same neuron, but also at the same synapse to self-modulate a highly stereotypic rhythmic behavior. A bidirectional glutamate-dependent inhibition system at this excitatory GABA synapse may increase the sensitivity and fidelity of this synapse to external modulators.

### UBR-1 regulates E/I signaling imbalance in the dual-release neuron

In *ubr-1* mutants, the activity and functional output of DVB is impaired. These defects are not caused by reduced GABA release or a development effect of *ubr-1*. Our studies demonstrate the critical source of defect is elevated glutamatergic signaling. We previously found that the whole-animal glutamate level is elevated in *ubr-1* mutants (Chitturi *et al*., 2018). We show here that in the DVB neuron, which exhibits a low level of inhibitory glutamatergic signaling, this leads to elevated inhibition of the defecation circuit activity.

It is difficult to demonstrate directly the co-release of neurotransmitters, but multiple lines of our experimental findings strongly support this idea. The vesicular glutamate transporter and the vesicular GABA transporter are co-expressed in the same neuron. Knock-down of EAT-4 in DVB fully rescues the defecation defects of *ubr-1* mutants. Thus, UBR-1 regulates the signaling strength between excitatory GABA and inhibitory glutamate transmission by gating glutamate content.

### Glutamate homeostasis and implications for the *JBS*

We identified here the consequence of changed synaptic E/I imbalance in absence of a functional UBR-1 protein due to disrupted glutamate homeostasis. Elevated glutamate, should it be ubiquitous cellular consequence of UBR1’s dysfunction, has interesting and important implication for the underlying cause of the *JBS*.

In the nervous system, altered E/I imbalance has been observed in multiple disorders including autism, Rett syndrome, mood disorders, and fragile X syndrome (Eichler & Meier, 2008; Gatto & Broadie, 2010; Marín, 2012; Zhang & Sun, 2011). Due to the disturbance of glutamate homeostasis, a change to the relative E/I signaling may contribute to neural development and functional impairment in the *JBS* patients. From this perspective, UBR1 might represent a potential regulator of the GABA/Glutamate signaling balance as well as glutamate metabolism.

## Materials and methods

### *C. elegans* strains and transgenic lines

*C. elegans* was grown at 22°C on the Nematode Growth Medium (NGM) plates with *E. coli* OP50 as a food source.(Brenner, 1974) Wild type animals are Bristol N2. *ubr-1(hp684)* was isolated from the EMS screen and other *ubr-1* alleles were generated using the CRISPR-Cas9 system (Chitturi *et al*., 2018). The other genetic mutants were obtained from the CGC (*Caenorhabditis Genetics Center*). The transgenic worms were created by microinjection according to standard protocols. Target DNA plasmids (∼50 ng/µL) were injected together with co-injections marker P*myo-2*::RFP or P*myo-3*::RFP at concentrations of 5-10 ng/µL. Typically, one transgenic line was tested in most of our rescue experiments except the *glc-3* behavioral rescue experiment, in which two lines of P*glc-3*::GLC-3 were examined. All strains used in this study are listed in Table S2.

### Defecation behavior assays

Defecation behavior was measured on NGM plates under standard conditions as previously described (Thomas, 1990). All behavioral tests were performed at 20-22°C. Worms were synchronized and well-fed young hermaphrodites adults were transferred to fresh NGM plates with OP50 for behavior assay during freely-moving state. The defecation phenotypes were scored under a Zeiss V16 microscope after 16-20 hours. Each animal was quantified in 10 minutes after the first pBoc step pass using the Etho program software (Liu & Thomas, 1994). Due to the aBoc step was difficult to distinguish between anterior body wall muscle contraction and pumping, we simplified the observation on the pBoc and the Exp steps (Iwasaki & Thomas, 1997). Exp/cycle was calculated as the ratio of the number of Exp/pBoc during a 10 min recording (Wang *et al*., 2013). The assay with GABA in *ubr-1* and *unc-25* mutants was performed as previously described (McLntire *et al*., 1993a).

### Molecular biology

All expression and rescue plasmids were generated via the multisite gateway system (Invitrogen, Thermo Fisher Scientific, Waltham, MA, USA) (Magnani *et al*, 2006). Three entry clones (Slot1, Slot2, Slot3) corresponding to promoter, target gene and fluorescent marker gene were recombined into pDEST™ R4-R3 Vector II via LR reaction, and then expression constructs were obtained. The genes *ubr-1*, *eat-4*, *glc-2*, *glc-3* and *glc-4* were cloned from wide-type genomic DNA. The *ubr-1* minigene plasmid contained a sequence of *unc-54-3’UTR*. The high fidelity polymerase Platinum HiFi Taq (Invitrogen) was used to amplify the cDNA sequence in the construction of the minigene. Plasmid sequence fidelity was verified by sequencing and enzyme digestion. The promoter lengths of P*ubr-1*, P*eat-4*, P*glc-2*, P*glc-3* and P*glc-4* were 1.6, 5.6, 2.0, 2.0 and 3.0 kb, respectively. All detailed information about plasmids and primers information are listed in Table S3-S4.

For cell-specific knockout experiments, we generate a *loxp* flanked (floxed) allele of *eat-4* minigene: P*eat-4*::loxP::EAT-4minigene(stop)::loxP::GFP (P*eat-4*::EAT-4^flox/flox^) that fully reverts the expulsion frequency of *ubr-1; eat-4* to *ubr-1* mutant. A Cre pilot strain was generated whose Cre recombinase is expressed in DVB neuron using the P*unc-47* promoter. Cre recognized *loxp* sequences and caused a site-specific deletion of *eat-4* DNA between two *loxp* sites.

### Fluorescence microscopy

The confocal images were obtained with a laser scanning confocal microscope (FV3000, Olympus) using 40x objectives (numerical aperture = 0.95). The neurons and muscles labeled with GFP or RFP have been imaged with excitation wavelength laser at 488 or 561 nm respectively. The worms were immobilized with 2.5 mM levamisole (Sigma-Aldrich) in M9 buffer. The images were processed and analyzed with ImageJ (National Institutes of Health).

### In vivo calcium imaging

The two integrated strains *hpIs468* (P*unc-47*::GCaMP6s::wCherry) and *hpIs582* (P*exp-1*::GCaMP6s::wCherry) were used for calcium imaging of DVB neuron and enteric muscle. Both strains exhibit normal defecation interval and Exp step. Young hermaphrodite adults were glued dorsally and imaged with a 60x water objective (numerical aperture = 1.0). In particular, animals are loosely immobilized with WormGlu (Histoacryl Blue, Braun, Germany) to a sylgard-coated cover glass covered with bath-solution (Sylgard 184, Dowcorning, USA) but leave the tail free in the recording solution. To keep the worms’ regular pumping and defecation activities, a small amount of OP50 (∼100 µl, OD_600_ = 1) was mixed into the recording solution, which consists of (in mM): NaCl 150; KCl 5; CaCl_2_ 5; MgCl_2_ 1; glucose 10; sucrose 5; HEPES 15, pH7.3 with NaOH, ∼330 mOsm.

Fluorescence images were acquired with excitation wavelength LED at 470 nm and a digital sCMOS camera at 100 ms per frame for 3 minutes. Data was collected from HCImage (Hamamatsu) and analyzed by Image Pro Plus 6.0 (Media Cybernetics, Inc., Rockville, MD, USA) and Image J (National Institutes of Health). For single-channel Ca^2+^ imaging, the GCaMP6s fluorescence intensity of the region of interest (ROI) was defined as *F*, the background intensity near the ROI was defined as *F_0_*. The true neuronal calcium fluorescence signal was obtained by subtracting the background signal from the ROI. *ΔF / F_0_ = (F − F_0_) / F_0_* was plotted over time as a fluorescence variation curve. For dual-channel Ca^2+^ imaging, with 470 nm and 590 nm excitation (LED, BioLED Light Source Control Module 9-24V DC), the emitted green (GCaMP6s) and red (wCherry) fluorescence were simultaneously collected under a wide-field microscope (Nikon LV-TV), which equipped with a dichroic beamsplitter (W-VIEW GEMINI, Japan), a sCMOS digital camera (Hamamatsu ORCA-Flash 4.0 V2) and a 60 x water objective (Nikon, Japan, numerical aperture = 1.0). And the ratiometric quantitation (*ΔR / R_0_*) was used to analyze the fluorescence change, in which *R* means the ratio of GFP/RFP (GCaMP6s/wCherry), *R_0_* means the minimal *R*, *ΔR* indicates (*R* − *R_0_*).

For extrinsic glutamate perfusion experiment in intact animals, glued worms were pretreated in the bath solution with 5.9 mM glutamate for 5 minutes, and then were imaged for another 3 minutes with 5.9 mM glutamate. For extrinsic glutamate perfusion experiment in dissected animals, DVB neuron was exposed by tail dissection as described for electrophysiological recording. To keep the circuit integrity, the dissection was performed just on the dorsal side in the tail near DVB soma. A glass capillary (2-6 MΩ) was used to punctured the epidermis and a small cut (5-10 μm width) was made to limit the possible damage or contents. Under this condition, almost all preparations exhibited regular rhythmic DVB Ca^2+^ activity (**Fig 7B**). Dissected DVB Ca^2+^ transient was recorded by the perfusion of 1 mM glutamate for ∼60 s and washing out by glutamate free bath solution.

### HEK293T expression and electrophysiology

GLC-3 expression in HEK293T cells: GLC-3 cDNA were flanked between SmaI and KpnI sites by PCR from cDNA of *C. elegans* and cloned into the expression vector pEGP-N1. HEK293T cells were cultured in the DMEM medium with 10% fetal bovine serum, 1% antibiotics (penicillin/streptomycin) at 5% CO_2_, and 37°C. After 24 hours of transfection of the plasmid with the liposome ExFect2000 (Vazyme, China), cells were patched using 4-6 MΩ resistant borosilicate pipettes (1B100F-4, World Precision Instruments, USA). Pipettes were pulled by micropipette puller P-1000 (Sutter, USA), and fire-polished by microforge MF-830 (Narishige, Japan). Membrane currents and I-V curve were recorded and plotted in the whole-cell configuration by pulse software with the EPC-9 amplifier (HEKA, Germany) and processed with the Igor Pro (WaveMetrics, USA) and Clampfit 10 software (Axon Instruments, Molecular Devices, USA). Membrane currents was recorded at −60 mV, I-V curve was recorded at a holding potential from −100 mV to +100 mV with 10 mV step voltages. Data were digitized at 10-20 kHz and filtered at 2.6 kHz. The pipette solution contained (in mM): KCl 140; MgCl_2_ 1; EGTA 10; HEPES 10; Na_2_ATP 4; pH 7.3 with KOH, ∼300 mOsm. The bath solution consisted of (in mM): NaCl 140; KCl 3.6; CaCl_2_ 2.5; MgCl_2_ 1; pH 7.3 with NaOH, ∼310 mOsm. For low extracellular Cl^−^ recording, the bath solution contained (in mM): Na-gluconate 140; KCl 3.6; CaCl_2_ 2.5; MgCl_2_ 1; pH 7.3 with NaOH, ∼310 mOsm. Chemicals were obtained from Sigma unless stated otherwise. Experiments were performed at room temperature (20-22°C).

### Statistical analysis and display

All defecation behavior data was displayed as bar graphs by GraphPad Prisim 8 (GraphPad Software Inc.). Calcium imaging data was mainly plotted using Matlab (MathWorks) for heat maps, Igor Pro (WaveMetrics) for curve graph and GraphPad for scatter plots, with each point representing a single worm test. The error bars represent the standard errors of the mean (SEM). An unpaired student’s *t*-test was used to analyze differences and calculate *p*-values when the comparison was limited to two groups. When multiple groups (more than two groups) of data were compared, ordinary one-way or two-way ANOVA analysis was performed for statistical analysis. Kolmogorov-Smirnov test was used to analyze amplitude cumulative fraction. The *p*-values is indicated as follows: ns, no significance, \**p* < 0.05, \*\**p* < 0.01, \*\*\**p* < 0.001, \*\*\*\**p* < 0.0001.

## Author Contributions

S.G., J.C., and M.Z. conceived experiments and S.G. wrote the manuscript. Y.L., J.C., B.Y., and Y.Z. performed experiments and analyzed data. J.W., T.P., W.H., and M.Z. contributed to the experiments. M.Z. edited the manuscript.

## Acknowledgements

We thank Gao lab members for comments and discussion. We thank Jun Meng for discussion, Ying Wang for technical assistance. This research was supported by the National Natural Science Foundation of China (32371189), the Major International (Regional) Joint Research Project (32020103007), the National Key Research and Development Program of China (2022YFA1206001), the National Natural Science Foundation of China (31871069), and the Canadian Institutes of Health Research (Foundation Scheme 154274), CSC Scholarship Program and the Overseas High-level Talents Introduction Program. We thank the Rey Interior (SPARC BioCentre, Sick Kids) for HPLC, *Caenorhabditis Genetics Center* for strains, which is funded by the NIH Office of Research Infrastructure Programs (P40 OD010440), for strains.

---

**Figure EV1.**
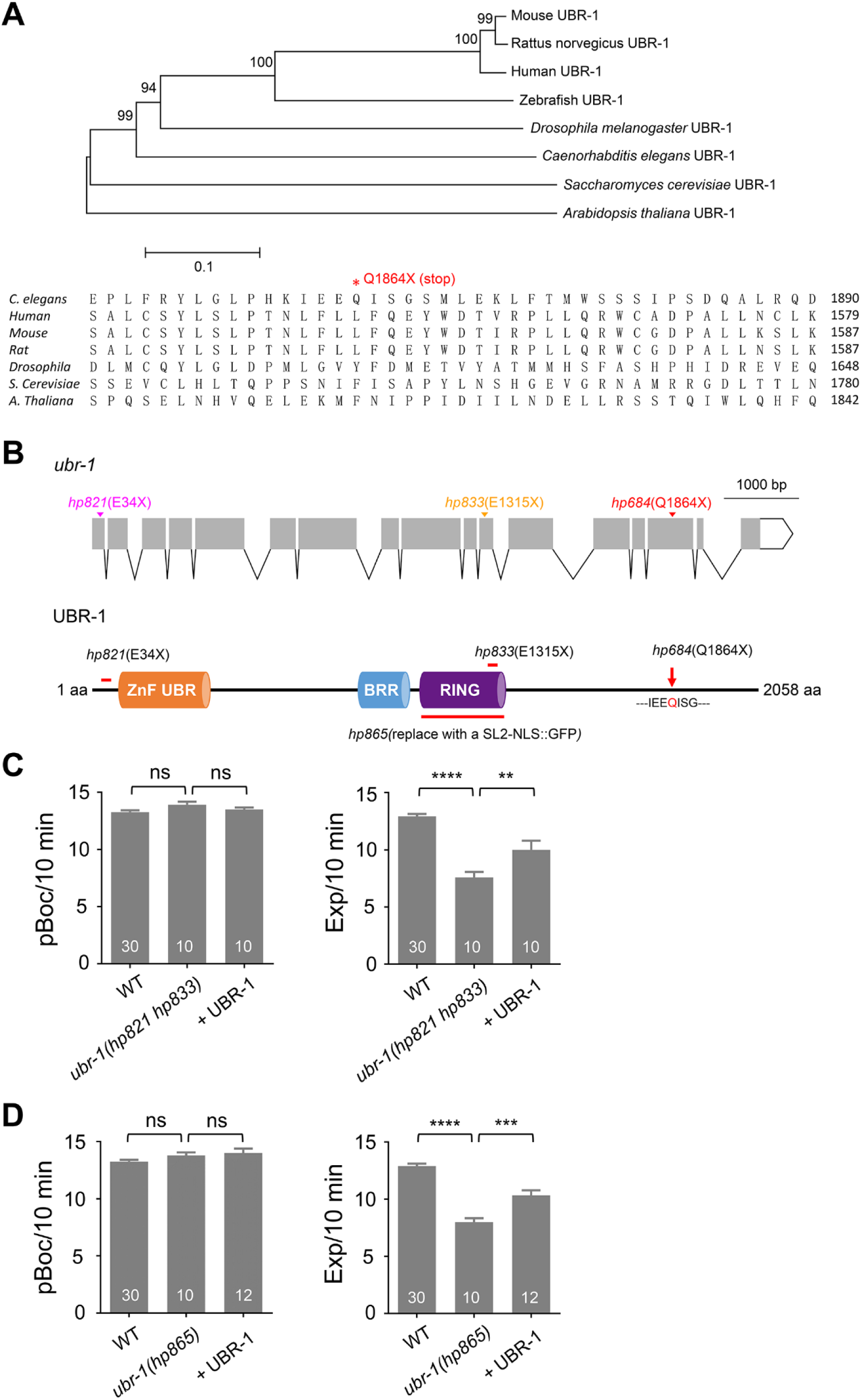
The *C. elegans* UBR-1 orthologs and structure. (A) Maximum likelihood tree of the phylogenetic relationship between UBR-1 orthologs from different species. Branch node labels likelihood ratio test values. (B) Structure of the *ubr-1* gene and UBR-1protein with significant mutants used in this study. Position and amino acid substitutions in *C. elegans* alleles are denoted. (C, D) *ubr-1* alleles exhibit same defecation defects. Quantification of frequency of pBoc and Exp in indicated genotypes. \*\**p* < 0.01; \*\*\**p* < 0.001; \*\*\*\**p* < 0.0001; one-way ANOVA test. n = 10-30 animals. Error bars, SEM.

**Figure EV2.**
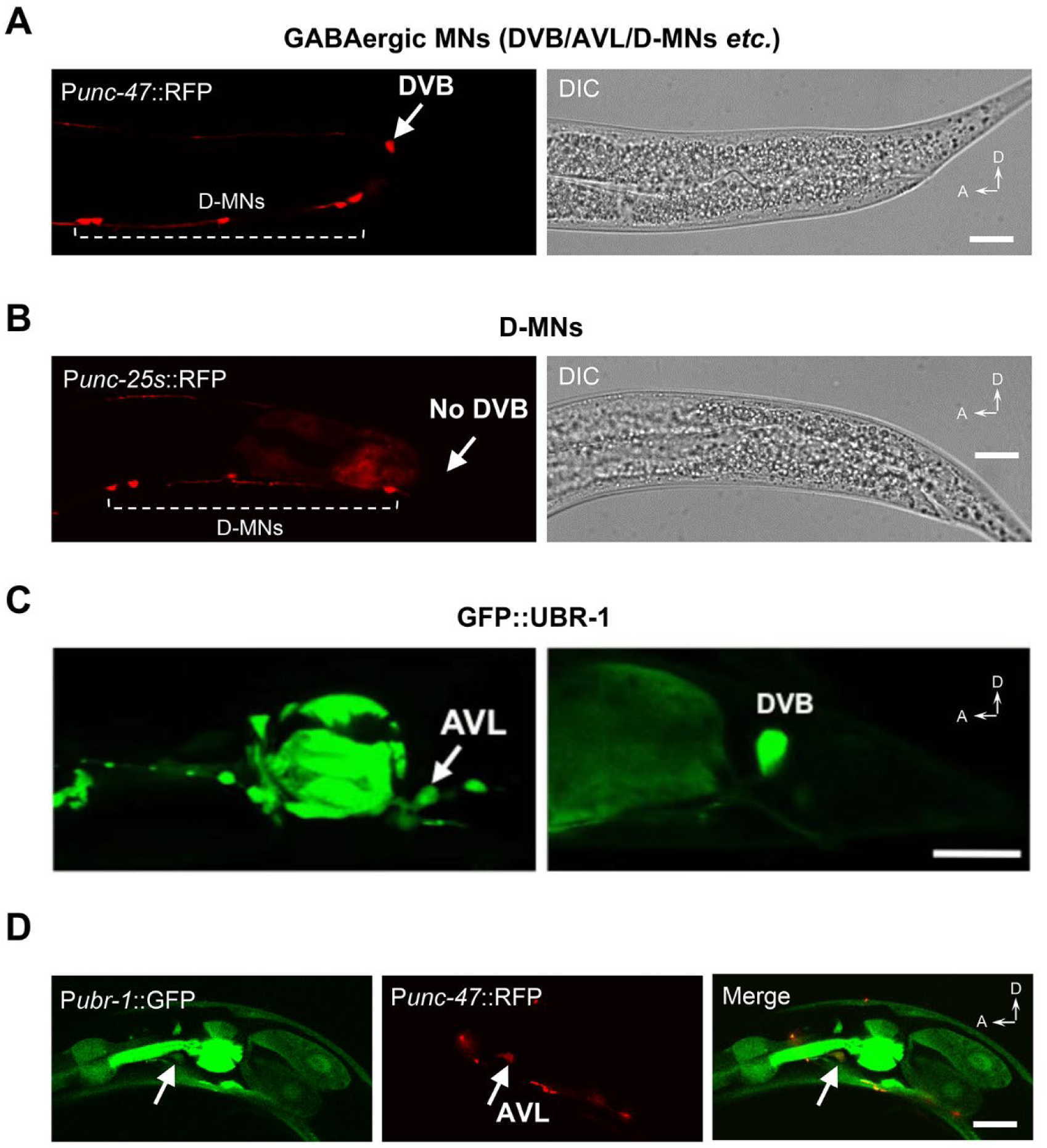
Expression pattern of *ubr-1* in AVL and DVB. (A) P*unc-47* driven RFP expresses in DVB and D-motor neurons. (B) Short fragment promoter (P*unc-25s*) used in this study show no DVB expression. (C, D) UBR-1 expression in AVL and DVB neurons. A anterior, D dorsal. Scale bar, 20 µm.

**Figure EV3.**
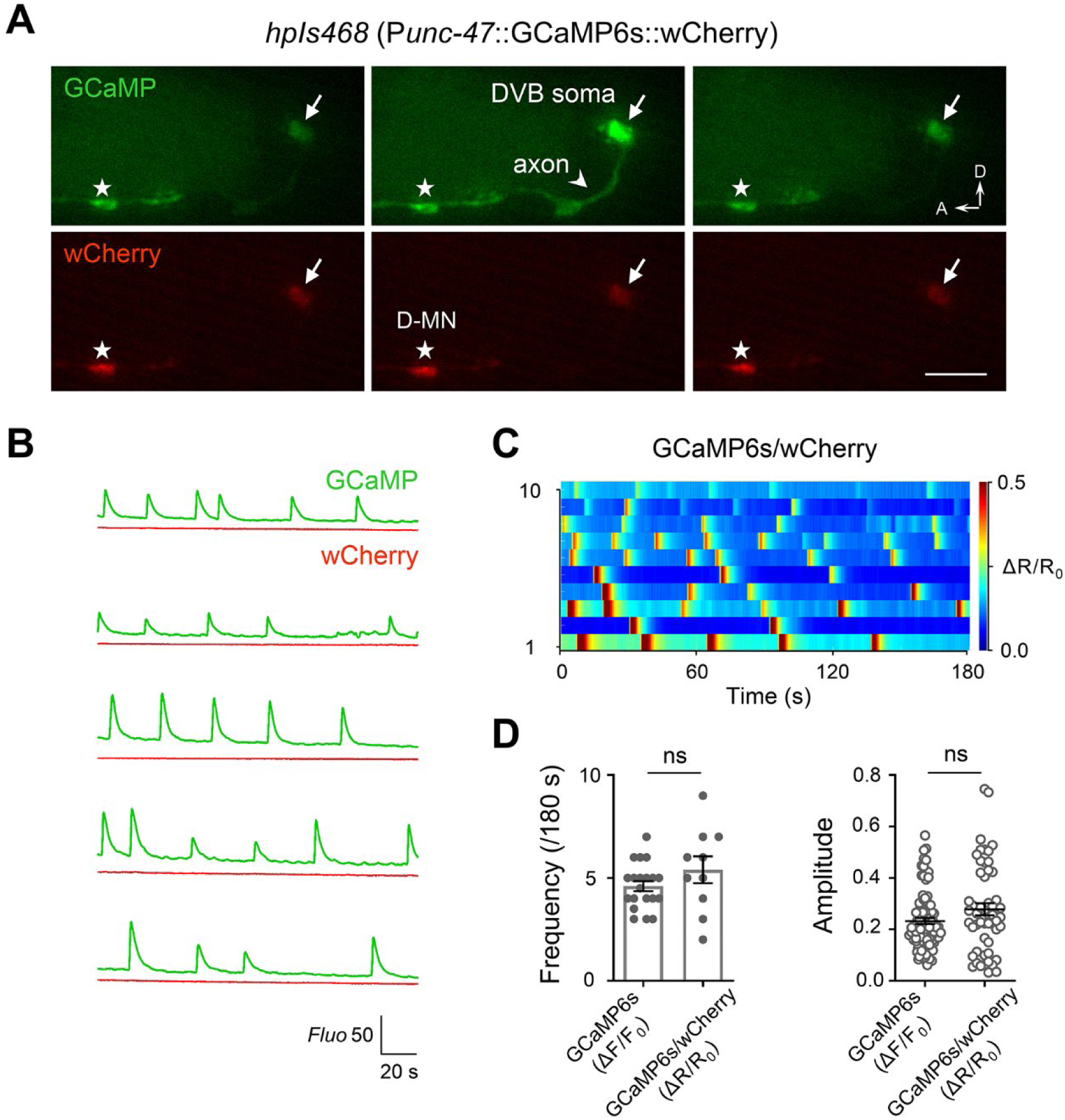
Dual-channel calcium imaging of DVB neuron. (A) Three real-time dual channel (GCaMP6s and wCherry) snapshots during a Ca^2+^ transient. The DVB soma was labeled by white arrows. Scale bar, 20 µm. (B) Five pairs of raw GFP and RFP traces were shown. Green and red traces denote the signals of GCaMP6s and wCherry, respectively. The wCherry fluorescence signals, as reference, show no change. (C) The color-map of the GCaMP6s/wCherry ratio. (D) Quantification and comparison of the frequency and single event peak amplitude between GCaMP6s and GCaMP6s/wCherry. n = 10-20 animals. ns, no significance; two-tailed unpaired *t*-test. A anterior, D dorsal. Error bars, SEM.

**Figure EV4.**
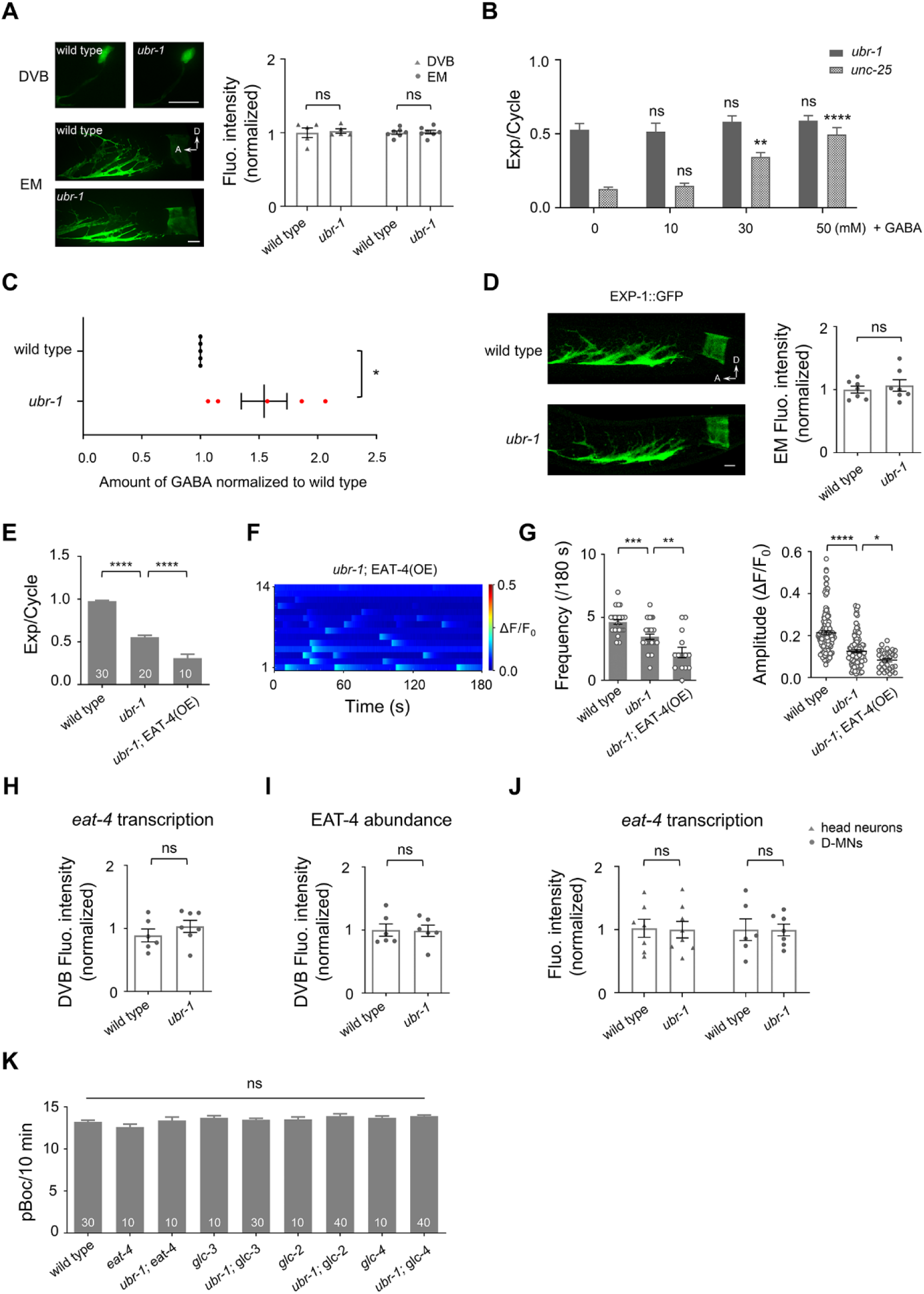
*ubr-1* does not affect the morphology of DVB and enteric muscles, as well as the expression of *eat-4*. (A) No obvious change in morphology and fluorescence intensity of DVB neuron (P*unc-47*::GFP) and enteric muscles (P*exp-1*::GFP) between wild type and *ubr-1* mutants. Scale bar, 10 µm. n = 5 animals. ns, no significance. (B) Exogenous addition of GABA (10-50 mM) to *ubr-1* and *unc-25* mutants. The expulsion defect of the *ubr-1* mutant was not improved after GABA application. (C) Free GABA were measured from whole worm lysates using HPLC. GABA level is significantly increased in *ubr-1* mutant compared to wild type animals. (D) The localization and expression density of the postsynaptic GABA receptor EXP-1 in wild type and *ubr-1* mutant. Scale bar, 20 µm. (E) Quantification of the expulsion rhythm in different genotypes. Overexpression of EAT-4 in GABAergic neurons reduces the expulsion frequency in *ubr-1* mutant. (F) Color maps summaries the DVB Ca^2+^ activity in EAT-4 overexpressed animals. (G) Quantification of the frequency and peak amplitude of DVB Ca^2+^ transients. *ubr-1* mutants exhibited significant reduced frequency and amplitude, which were further decreased by EAT-4 overexpression in GABAergic neurons. (H-J) No obvious change in the fluorescence levels of DVB neuron between wild type and *ubr-1* mutant, as well as head neurons and a subset of D-MNs. (K) Quantification of pBoc frequency in indicated genotypes. \*\**p* < 0.01; \*\*\*\**p* < 0.0001; two-way ANOVA test in A and B. \**p* < 0.05; two-tailed unpaired *t*-test in C and D, H-J. \**p* < 0.05; \*\**p* < 0.01; \*\*\**p* < 0.001; \*\*\*\**p* < 0.0001; one-way ANOVA test in E, G and K. A anterior, D dorsal. Error bars, SEM.

**Figure EV5.**
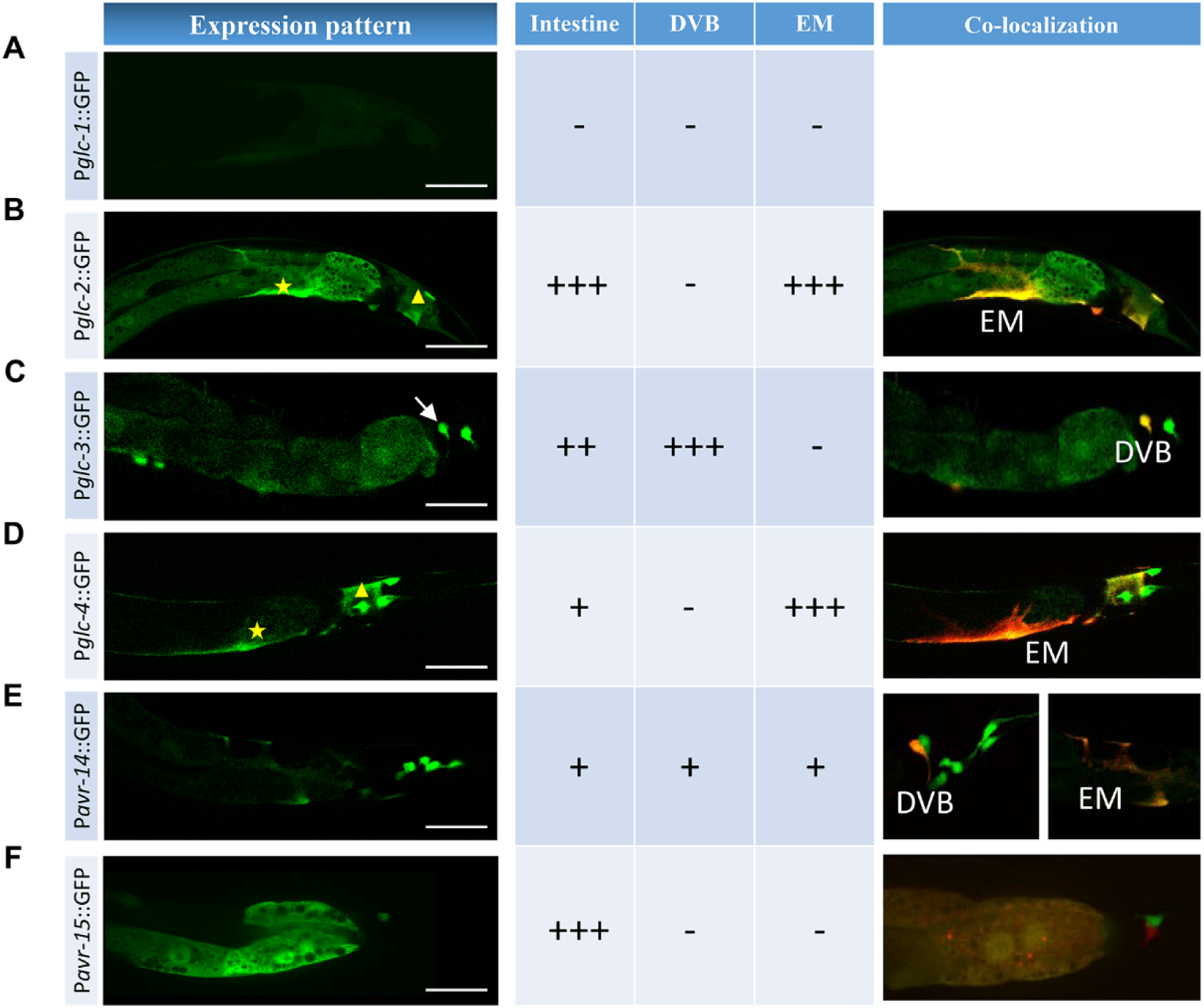
Expression pattern of glutamate-gated chloride channels in the tail. (A-F) The expression pattern of six GluCl genes from *glc-1* to *glc-4*, as well as *avr-14* and *avr-15*. −, no expression; +, slight expression; ++, moderate rescue; +++, strong expression. Scale bar, 20 µm. Yellow stars and triangles label the enteric muscle and anal depressor muscle, respectively. White arrows label the DVB neuron.

